# Sequence type diversity amongst antibiotic-resistant bacterial strains is lower than amongst antibiotic-susceptible strains

**DOI:** 10.1101/2022.11.23.517742

**Authors:** Anjani Pradhananga, Lorena Benitez-Rivera, Candace Clark, Kaho Tisthammer, Pleuni Simone Pennings

## Abstract

The increasing number of antibiotic resistant bacterial infections is a global threat to human health. Antibiotic resistant bacterial strains generally evolve from susceptible strains by either horizontal gene transfer or chromosomal mutations. After evolving within a host, such resistant strains can be transmitted to other hosts and increase in frequency in the population at large. Population genetic theory postulates that the increase in frequency of an adaptive trait can lead to signatures of selective sweeps. One would thus expect to observe reduced genetic diversity amongst that part of the population that carries the adaptive trait. Specifically, if the evolution of new resistant strains is rare, it is expected that resistant strains represent only a subset of the diversity of susceptible strains. It is currently unknown if diversity of resistant strains is indeed lower than diversity of susceptible strains when considering antibiotic resistance. Here we show that in several bacterial species in several different datasets, sequence-type diversity amongst antibiotic-resistant bacterial strains is indeed lower than amongst antibiotic-susceptible strains in most cases. We re-analysed eight existing clinical datasets with *Escherichia coli, Staphylococcus aureus* and *Enterococcus faecium* samples. These datasets consisted of 53 - 1094 patient samples, with multi-locus sequence types and antibiotic resistance phenotypes for 3 - 19 different antibiotics. Out of 59 comparisons, we found that resistant strains were significantly less diverse than susceptible strains in 51 cases (86%). In addition, we show that sequence-type diversity of antibiotic-resistant strains is lower if resistance is rare, compared to when resistance is common, which is consistent with rare resistance being due to fewer evolutionary origins. Our results show that for several different bacterial species, we observe reduced diversity of resistant strains, which is consistent with the evolution of resistance driven by selective sweeps stemming from a limited number of evolutionary origins. In future studies, more detailed analysis of such sweep signatures is warranted.

## Introduction

Antimicrobials are drugs that help us combat diseases caused by pathogens that infect some bacteria, viruses, parasites, and fungi. An increase of antimicrobial resistance threatens public health and well-being. According to the World Bank, an additional 24 million people will be added into the extreme poverty category by 2030 (Jonas et al., 2017), as an impact of antimicrobial resistance. In addition, antibiotic resistance could cause 10 million deaths by 2050, and also result in the loss of 100 trillion USD worldwide (Founou et al., 2017). In the US alone there are estimated to be 2.8 million antibiotic-resistant infections and 35,000 deaths each year (CDC, 2020) For all these reasons, antibiotic-resistant bacterial infections are receiving a lot of attention (Balsalobre et al., 2014, Barbosa & Levy, 2000, and Pendleton et al., 2013).

While it is clear that antibiotic resistance is a big problem worldwide, the population genetics of drug resistance evolution outside of the laboratory is not well understood. If we knew more about the origins of antibiotic resistant bacteria and how they spread, this would likely help us prevent antibiotic-resistant infections. Several important studies have determined the origins of drug resistant strains. For example, Enright and colleagues showed that *Staphylococcus aureus* has acquired the SCC*mec* element at least 11 times (Enright et al., 2002). Another study focused only on sequence type 5 of *S. aureus* and showed that the SCC*mec* element was imported into ST5 *S. aureus* strains at least 23 times (Nubel et al., 2008). The same study also reported that most of the resulting MRSA strains remain local in one or a few countries (Nubel et al., 2008). In addition to multiple origins of resistant strains, it has been shown that in most cases, susceptible strains co-exist with resistant strains (Austin et al., 1999; Blanquart et al., 2017). These results suggest that antibiotic resistance evolution may be best described by local and incomplete soft sweeps from multiple origins.

If the number of different origins of antibiotic resistant strains is fairly small, one may expect that resistant strains are overall less diverse and have fewer sequence types than susceptible strains. However, if the number of different origins of antibiotic resistant strains is large, as suggested by (Nubel et al., 2008), diversity of resistant strains may be as high as diversity of susceptible strains. We set out to test for different bacterial species and different antibiotics, whether resistant strains are indeed less diverse than susceptible strains.

In this paper we analyze 38 antibiotics found in six different published studies (eight datasets) focusing on three pathogens: *E. coli, S. aureus* and *E. faecium*. For each study and each antibiotic, we determine whether the resistant strains show lower sequence type diversity compared to the susceptible strains as measured by the Gini-Simpson Index, the Inverse Simpson Index, and the Shannon’s Diversity Index (H’) (Jost, 2006; Morris et al., 2014; Simpson, 1949).

With the collected data, we test two hypotheses. First, we determine whether resistant strains are less diverse than susceptible strains. Second, we determine whether diversity of resistant strains is affected by how rare the specific resistance is.

## Methods

### Data collection

We used data from a convenience sample of published papers that reported antibiotic resistance and multi-locus sequence typing (MLST) for individual samples of bacterial infections. The published papers were found through the NCBI, San Francisco State University (SFSU), and Google scholar databases. The main criteria for inclusion of the data was that the paper include both resistant and susceptible strains for each of the antibiotics, mentions all the antibiotics used, and reported MLST information (sequence types) for each patient sample. Papers were excluded if they only reported data for resistant samples or if they only reported summary statistics, but not the information for each sample.

Each dataset we collected contains a list of samples (one sample per patient), and for each sample we know whether it is resistant or susceptible to a list of antibiotics. For each drug, we can therefore split a dataset in a population of resistant and susceptible samples. Because we also know the multi-locus sequence typing for each sample, we can calculate the sequence type diversity of the population of resistant samples and the sequence type diversity of the population of susceptible samples (using different measures of diversity). The focus of the study is to determine whether the resistant populations are less diverse than the susceptible populations.

Multi-locus sequence typing (MLST) analysis was used to calculate diversity. Multi-locus sequence types are commonly used to characterize bacterial strains based on the sequence of a small number of standard housekeeping genes. A combination of each unique allele of the housekeeping genes is assigned a ST number such as ST3, ST5, ST15, ST131 and so on (Maiden et al., 1998; Adiri et al., 2003; Enright et al., 2000; and Homan et al., 2002).

In total we collected data from six published papers. Three of these papers reported data on *E. coli* infections: Yamaji et al., 2018, Adams-Sapper et al., 2013, and Kallonen et al., 2017. Two papers are based on the study of *S. aureus* infections Wurster et al., 2018 and Manara et al., 2018. One paper is based on *E. faecium* infections Galloway-Peña et al., 2009. The Yamaji *E. coli* article includes two datasets which are collected from the same location, but 17 years apart, therefore we consider it as two different datasets for our analysis. Furthermore, the Kallonen *E. coli* article includes two datasets, collected from national and local sites; which we consider as two different datasets for our analysis. Therefore, in total eight different datasets from the six published papers were collected for the analysis (Table 1)

**Table 1:**
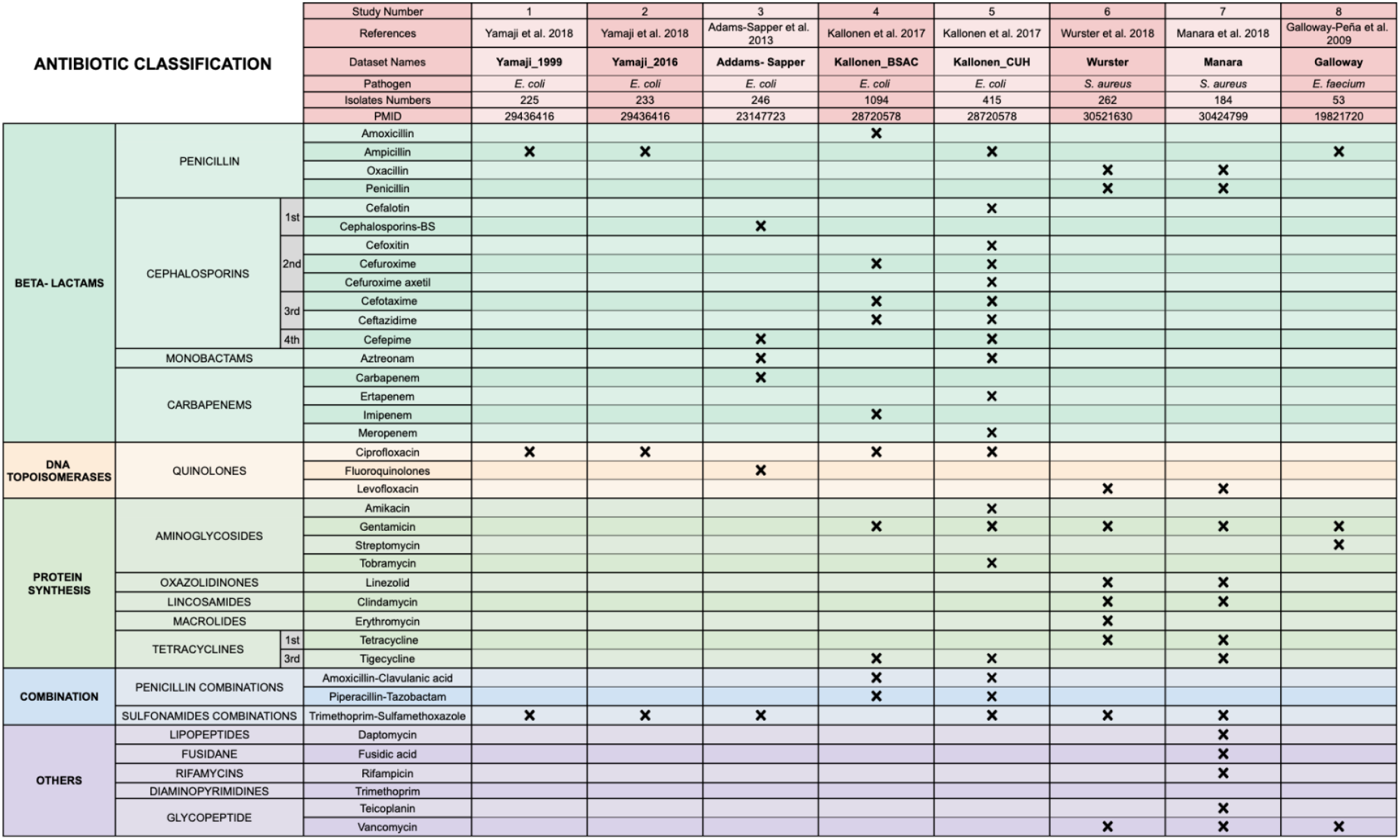
An overview of all the datasets included in the study with information on the antibiotics that are investigated in each dataset.

#### *Escherichia coli (E. coli)* datasets

For *E. coli*, there are five datasets from three different published papers:

1. **Yamaji_1999** consists of **225 isolates** from patients with urinary tract infections (UTIs) collected between 1999 and 2000 at the University of California, Berkeley, California, USA. Resistance phenotypes are available for **three different antibiotics** (ampicillin, ciprofloxacin and trimethoprim-sulfamethoxazole).
2. **Yamaji_2016** consists of **233 isolates** from patients with urinary tract infections (UTIs) collected between 2016 and 2017 at the University of California, Berkeley, California, USA. Resistance phenotypes are available for **three different antibiotics** (ampicillin, ciprofloxacin and trimethoprim-sulfamethoxazole).
3. **Addams-Sapper** consists of **246 isolates** from patients with bloodstream infections collected between 2007 and 2010 at San Francisco General Hospital (SFGH), California, USA. Resistance phenotypes are available for **six different antibiotics** (cefepime, aztreonam, cephalosporins-BS, fluoroquinolones, and trimethoprim-sulfamethoxazole).
4. **Kallonen_BSAC** consists of **1094 isolates** from patients with bacteremia collected between the years of 2001 and 2011 by the British Society for Antimicrobial Chemotherapy (BSAC) from 11 hospitals across England. Resistance phenotypes are available for **10 different antibiotics** (amoxicillin, amoxicillin-clavulanic acid, cefotaxime, ceftazidime, cefuroxime, ciprofloxacin, gentamicin, imipenem, piperacillin-tazobactam, and tigecycline).
5. **Kallonen_CUH** consists of **415 isolates** from patients with bacteremia collected between the years of 2006 and 2012 from the local diagnostic laboratory at the Cambridge University Hospitals in England. Resistance phenotypes are available for **19 antibiotics** (amikacin, amoxicillin-clavulanic acid, ampicillin, aztreonam, cefalotin, cefepime, cefotaxime, cefoxitin, ceftazidime, cefuroxime, cefuroxime axetil, ciprofloxacin, ertapenem, gentamicin, meropenem, piperacillin-tazobactam, tigecycline, tobramycin, and trimethoprim)

#### *Staphylococcus aureus (S. aureus)* datasets

There were two different datasets for *S. aureus* from two different published papers:

5. **Wurster** consists of **262 clinical isolates** from ocular and otolaryngology infections collected from January to December 2014 from Massachusetts Eye and Ear, a Harvard teaching hospital, Boston, Massachusetts, USA. Resistance phenotypes are available for **11 antibiotics (**clindamycin, erythromycin, gentamicin, levofloxacin, linezolid, oxacillin, penicillin, tetracycline, trimethoprim-sulfamethoxazole, and vancomycin)
6. **Manara** consists of **184 clinical isolates** from respiratory tract infections (RTIs), soft tissue and skin lesions of patients collected from 2013 to 2015 at the Anne Meyer’s Children’s University Hospital, Florence, Italy. Resistance phenotypes are available for **14 antibiotics (**clindamycin, daptomycin, fusidic acid, gentamicin, levofloxacin, linezolid, oxacillin, penicillin, rifampicin, teicoplanin, tetracycline, tigecycline, trimethoprim-sulfamethoxazole, and vancomycin)

#### *Enterococcus faecium (E. faecium)* dataset

There is one dataset with *Enterococcus faecium* samples:

7. **Galloway** consists of **53 clinical isolates** from nosocomial patients collected from 1971 to 1994 at diverse geographic locations in the United States. Resistance phenotypes are available for **4 antibiotics** (ampicillin, gentamicin, streptomycin, and vancomycin)

### Data preparation

Custom R scripts were used to prepare the data from the different sources for analysis. Specifically, for each dataset, we created a.csv file which contains information of each sequence type, each drug, the number of resistant samples and the number or susceptible samples. Before analysis of the data, we removed any sequence types that were marked as “minor”, “ND” or “-” because these categories could include multiple different sequence types. Table 2 shows an example dataset (the smallest of the datasets) for illustration.

**Table 2.** Example dataset. We show here Yamaji_1999, which is the smallest of the datasets.

| Sequence Type | Drug | Number of resistant samples | Number of susceptible samples |
| --- | --- | --- | --- |
| ST 95 | Ampicillin | 1 | 33 |
| ST 127 | Ampicillin | 3 | 21 |
| ST 73 | Ampicillin | 6 | 17 |
| ST 69 | Ampicillin | 19 | 7 |
| ST 131 | Ampicillin | 2 | 5 |
| ST 10 | Ampicillin | 2 | 9 |
| ST 95 | Trim-Sulf | 1 | 33 |
| ST 127 | Trim-Sulf | 1 | 23 |
| ST 73 | Trim-Sulf | 1 | 22 |
| ST 69 | Trim-Sulf | 17 | 9 |
| ST 131 | Trim-Sulf | 1 | 6 |
| ST 10 | Trim-Sulf | 2 | 9 |

## Data accessibility

All the data and R scripts are available on Github (Antibiotic_Resistance_Data_Analysis.git>).

## Data analysis

This study is based on eight different datasets from six different published papers. The main analysis is to compare resistant and susceptible sequence types (ST) against the antibiotics used in each sample. Because antibiotics resistance is a newly evolved trait for the bacterial species, the expectation is that sequence type diversity is lower among resistant populations when they are compared to the susceptible ones.

Several approaches for the analysis of the datasets to study the antibiotic diversity among resistant and susceptible populations were taken. First, calculation of diversity by using Gini-Simpson Index, Inverse Simpson Index and Shannon’s Diversity Index (H’). The three indices were applied to the resistant and susceptible populations (all resistant, and susceptible samples) for each antibiotic in each dataset. Second, analysis of significance by using bootstrapping to determine whether the resistant populations were significantly less diverse than the susceptible populations. Third, analysis of observed levels of diversity by a linear model to determine which characteristics of each dataset can explain the level of diversity.

### Calculation of diversity for resistant and susceptible antibiotic populations

We used three common measures of diversity: Gini-Simpson Index, Inverse Simpson Index, and Shannon’s Diversity Index (H’) The Gini-Simpson Index and the Inverse Simpson Index are popular indices to measure diversity and they are often used to quantify biodiversity. The Gini-Simpson Index is calculated by this formula: 1- ∑(p_i_)^2^ where p_i_ is the proportional abundance of the *i*th sequence type. In the Gini-Simpson Index the values range from zero to one, where 0 represents no diversity and 1 represents the highest diversity. The Inverse Simpson Index is calculated by: 1 / ∑(p_i_)^2^ and the index values can be higher than one. Another popular diversity index is the Shannon’s Diversity Index (H’), which is calculated as - ∑p_i_ ln(p_i_). Shannon’s Diversity Index measures the uncertainty in predicting the sequence type of the identity of sequence type samples that are randomly taken from the datasets. Shannon’s Diversity Index values can be higher than one. For the three measures of diversity indices, the higher the indices values, the higher the diversity for the resistant or susceptible population (Jost, 2006; Morris et al., 2014; Simpson, 1949).

### Analysis of significance for resistant and susceptible antibiotic populations

After calculating the diversity of the resistant populations versus the susceptible populations for each drug in each datasets, a bootstrapping approach was considered to test our hypothesis in order to determine level of significance (Kulesa et al., 2015).

For each antibiotic in each dataset, a randomized set of resistant and susceptible populations were simulated 1000 times by resampling with replacement. Diversity of resistant and susceptible populations were determined by calculating diversity values for these randomized groups. In each simulation the total number of resistant and susceptible samples remained the same as that of collected data. We then determined whether the observed diversity values were more extreme than the 95% interval of the simulated data.

### Regression analysis: what determines diversity

The main focus of the study is to understand better which factors determine diversity of resistant populations. While our dataset is not extensive enough to test for many different factors, a linear model was used to determine whether the fraction of resistant populations (i.e., what fraction of patients in a dataset carried resistance to a specific antibiotic) has an effect on observed diversity. However, in order to do this, we have to use a normalized version of the Gini-Simpson Index (GSI): GSI_nor = GSI / GSI_max.

When there are only a few patients with resistant strains, the diversity cannot be as high as when there are many patients with resistant strains. For example, if there are just two patients with resistant strains, we can have at most two sequence types, with each a frequency of 50%. In that case the maximum possible value of the Gini-Simpson Index is 1-(0.5^2 +0.5^2) = 0.5. Therefore, even if resistant and susceptible labels were randomly assigned to the patients, the findings of the fraction of patients with resistant strains would still show an effect on diversity. To deal with this issue, we decided to use a normalized version of Gini-Simpson Index by dividing the observed Gini-Simpson Index value by the maximum possible value of the Gini-Simpson Index given the sample size, similar to an approach taken by (Garud & Rosenberg, 2015). We find that, as we hoped, when we randomly assign labels (resistant and susceptible), the fraction of patients with resistant strains has no effect on the normalized Gini-Simpson Index values in simulated data.

To determine if the fraction of patients with resistant strains does have an effect on the actual observed normalized Gini-Simpson Index (GSI) values, we used a generalized linear model in R using the following formula: glm(formula = GSI_nor ∼ FracRes + Dataset, data = Data).

## Results

### Diversity among susceptible and resistant Sequence Types (STs): 3 examples

For each of the antibiotics in each of the datasets we calculated the diversity for resistant and susceptible strains using the Gini-Simpson Index. We used a bootstrapping approach to determine whether resistant samples were significantly less diverse than susceptible samples. By comparing two of the datasets Wurster and Kallonen_CUH we show whether the diversity of the penicillin, oxacillin and tobramycin between susceptible and resistant strains are similar or different. For penicillin in the Wuster dataset, the Gini-Simpson Index bars (Figure 1A) represent the penicillin-susceptible (teal) and penicillin-resistant (red) strains. The Gini-Simpson Index values for penicillin-susceptible strains is 0.90, which is nearly the same as the penicillin-resistant strains having a value of 0.91. Penicillin in the Wuster dataset has non-significant diversity (p > 0.5) between susceptible and resistant strains based on the Gini-Simpson Index values. In addition, the pie charts (Figures 1B and 1C) show similar diversity in sequence strains in both penicillin-resistant and penicillin-susceptible. However, for oxacillin in the Wuster dataset (Figure 1D) oxacillin-susceptible (teal) and oxacillin-resistant (red) show a clear difference between susceptible and resistant strains. The Gini-Simpson Index for oxacillin-susceptible strains is 0.96 and oxacillin-resistant strains is 0.73, with highly significant diversity (p<0.001 using a permutation test). The pie charts (Figure 1E and 1F) show that the oxacillin-resistant strains are visibly less diverse. In the case of tobramycin for the Kallonen_CUH dataset (Figure 1G), tobramycin-susceptible (teal) and tobramycin-resistant (red) strains show highly significant diversity (p<0.001). The Gini-Simpson Index values for tobramycin-susceptible is 0.93 and tobramycin-resistant is 0.47, which are significantly different. The pie charts show more diversity for tobramycin-susceptible strains (Figure 1H) and clearly show less diversity for tobramycin-resistant strains (Figure 1I).

**Figure 1.**
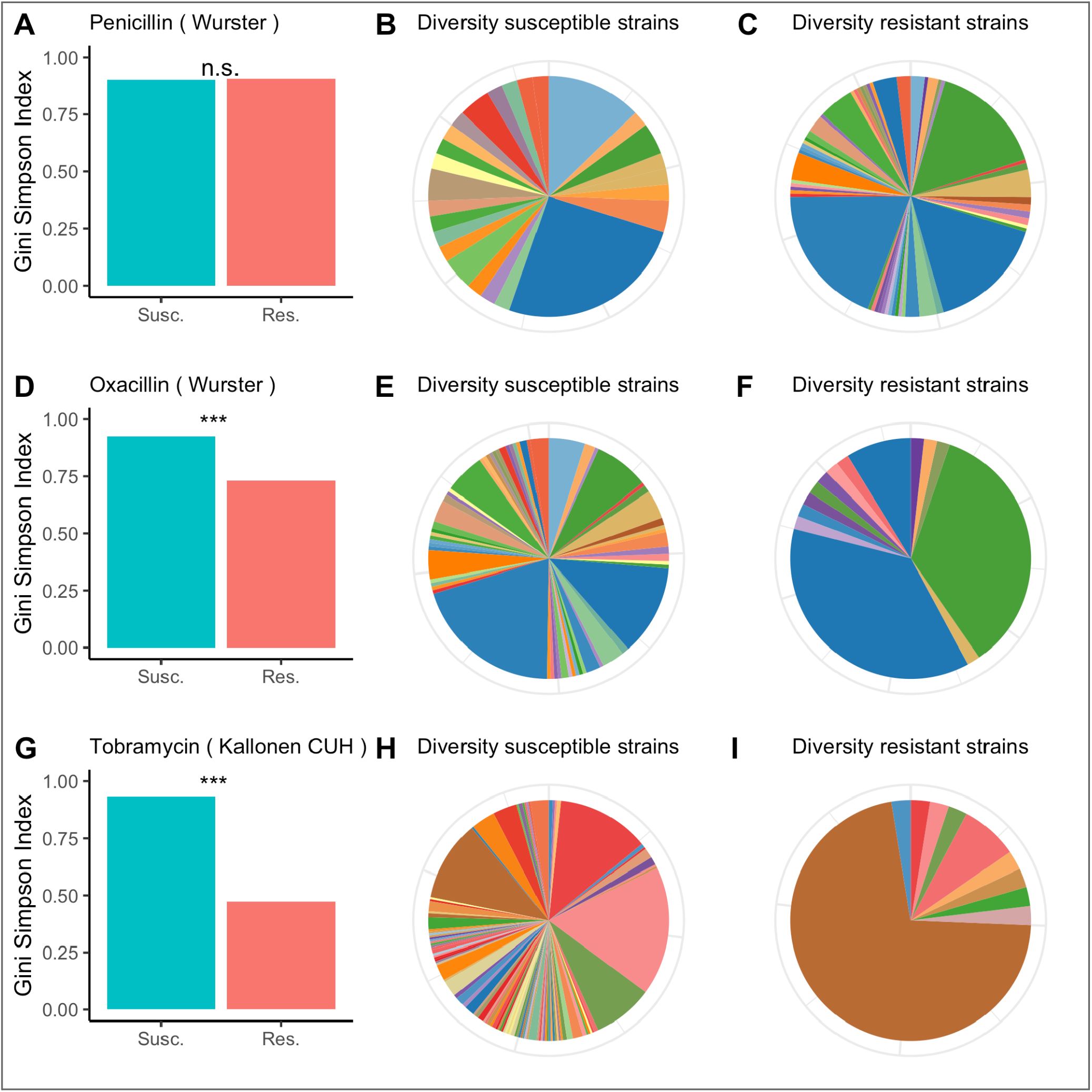
Diversity among susceptible and resistant populations. A. Gini-Simpson Index of diversity among penicillin-susceptible samples (teal) and penicillin-resistant samples (red) for the Wurster dataset. B. Sequence type abundance among penicillin-susceptible samples in the Wurster dataset. In the pie charts, only sequence types with abundances higher than 5% are labeled. C. Sequence type abundance among penicillin-resistant samples in the Wurster dataset. D, E, F: like A, B, C but for oxacillin-susceptible and oxacillin-resistant samples in the Wurster dataset. G, H, I: like A, B, C but for tobramycin-susceptible and tobramycin-resistant samples in the Kallonen_CUH dataset.

Plots for other antibiotics and all datasets are available in the github repository.

### Diversity values for resistant samples are usually lower than for susceptible samples

We calculated three diversity indices for all datasets, but will focus here on the Gini-Simpson Index for diversity. We plotted the Gini-Simpson Index for resistant and susceptible strains for each drug for each of the datasets (Figure 2A) and the fraction of samples that was found to be resistant (Figure 2B). As can be seen in the figure, in most of the cases, resistant strains were significantly less diverse than resistant strains.

**Figure 2.**
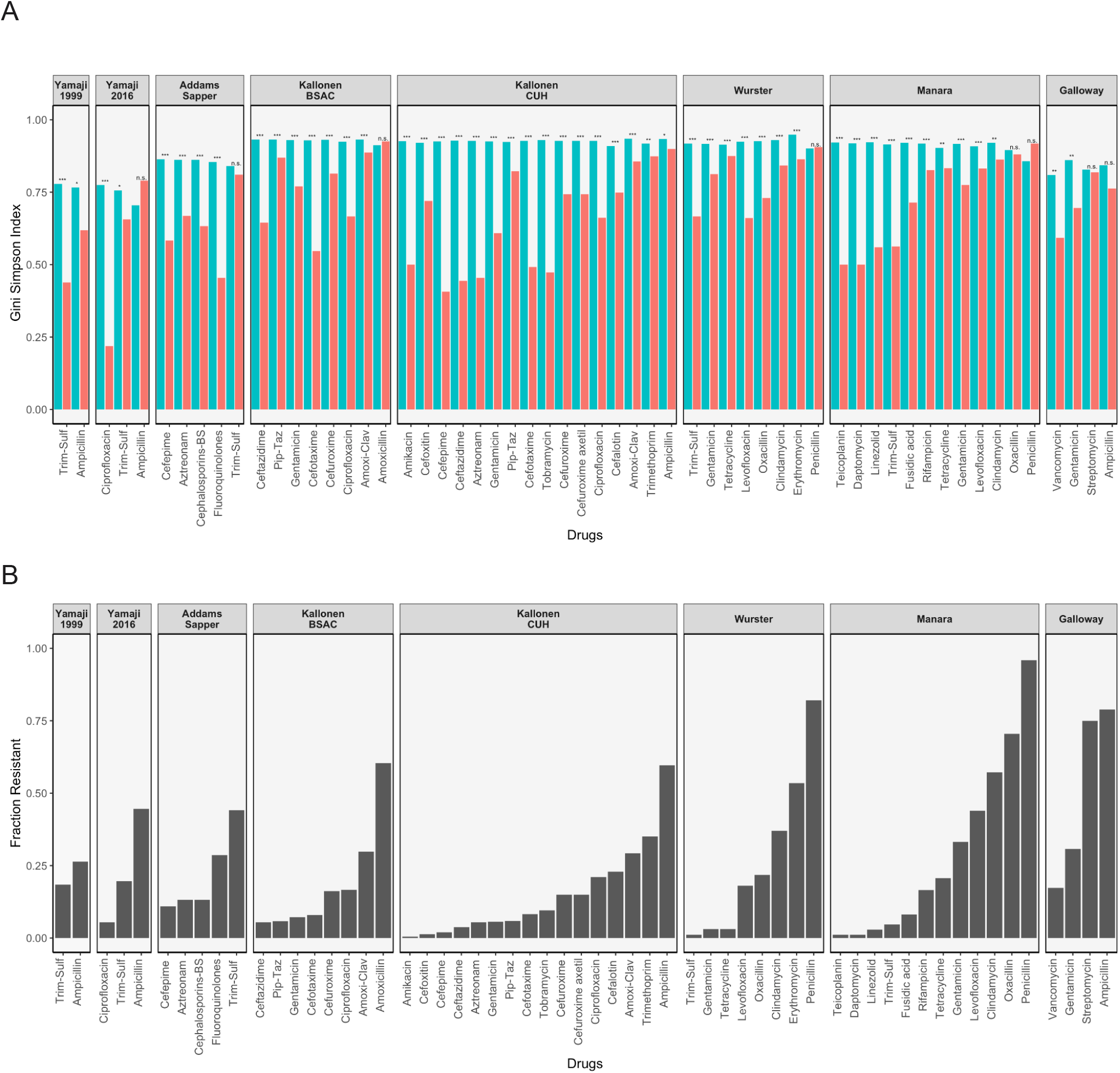
Diversity values of resistant samples. A: Comparison of the Gini-Simpson Index for resistant and susceptible strains in all datasets. Trim-Sulf (trimethoprim-sulfamethoxazole), Pip-Taz (piperacillin-tazobactam), Amoxi-Clav (amoxicillin-clavulanic acid). B: Resistant fraction ratio.

#### *E. coli* datasets

In the Yamaji dataset from 1999-2000, the samples were tested against three antibiotics, but no samples were resistant against ciprofloxacin, so it was not possible to calculate diversity. For ampicillin and trimethoprim-sulfamethoxazole, the resistant strains were significantly less diverse than the susceptible strains, although the difference was bigger and more significant for trimethoprim-sulfamethoxazole. In the later data (2016-2017) from the same study, we see that resistance has gone up for all three drugs (Figure 2B). In terms of diversity, we see the lowest diversity for the (still rare) ciprofloxacin resistant strains, higher diversity for the more common trimethoprim-sulfamethoxazole resistant strains and even higher diversity for the very common ampicillin-resistant strains. In the latter case, the resistant strains even display more diversity than the susceptible strains.

In the Addams-Sapper dataset resistance was observed against 5 drugs (cefepime, aztreonam, cephalosporins-BS, fluoroquinolones and trimethoprim-sulfamethoxazole). In all cases except for trimethoprim-sulfamethoxazole, the resistant strains were significantly less diverse than the susceptible strains (Figure 2A). Trimethoprim-sulfamethoxazole was the most common resistance in this dataset (Figure 2B).

In the Kallonen-BSAC dataset, resistance was observed against 7 drugs (ceftazidime, gentamicin, cefotaxime, cefuroxime, amoxicillin-clavulanic acid and amoxicillin). In all cases except for amoxicillin, the resistant strains were significantly less diverse than the susceptible strains (Figure 2A) and amoxicillin was the most common resistance in this dataset. In the second Kallonen dataset (Kallonen-CUH), resistance was observed against 15 drugs (amikacin, cefoxitin, cefepime, ceftazidime, aztreonam, gentamicin, piperacillin-tazobactam, cefotaxime, tobramycin, cefuroxime, cefuroxime axetil, ciprofloxacin, cefalotin, amoxicillin-clavulanic acid and ampicillin). In all cases, resistant strains were significantly less diverse than the susceptible strains (Figure 2).

#### *S. aureus* datasets

In the Wurster dataset, resistance was observed against 8 drugs (trimethoprim-sulfamethoxazole, gentamicin, tetracycline, levofloxacin, oxacillin, clindamycin, erythromycin and penicillin). In all cases except penicillin, resistant strains were significantly less diverse than the susceptible strains (Figure 2A). Penicillin resistance was the most common resistance in this dataset (Figure 2B). In the Manara dataset, resistance was observed against 12 drugs (teicoplanin, daptomycin, linezolid, trimethoprim-sulfamethoxazole, fusidic acid, rifampicin, tetracycline, gentamicin, levofloxacin, clindamycin, oxacillin, penicillin). In all cases except oxacillin and penicillin, resistant strains were significantly less diverse than the susceptible strains (Figure 2). Oxacillin and penicillin resistance were the most common resistance in this dataset.

#### *E. faecium* dataset

In the Galloway-Peña dataset, resistance was observed against 4 drugs (vancomycin, gentamicin, streptomycin and ampicillin). In two cases (vancomycin and gentamicin) resistant strains were significantly less diverse than the susceptible strains (Figure 2). In the other two cases (streptomycin and ampicillin) for which drug resistance was more common, there was no significant difference in diversity between resistant and susceptible strains.

Out of 59 comparisons of the Gini-Simpson Index between resistant and susceptible strains, in 51 cases we found that resistant strains were significantly less diverse. In 4 cases, there was a small difference, but it was not significant and in 4 other cases resistant strains had higher diversity than susceptible strains: ampicillin (*E. coli* in the Yamaji 2016 dataset), amoxicillin (*E. coli* in the Kallonen BSAC dataset), penicillin (*S. aureus* in the Wurster dataset) and penicillin (*S. aureus* in the Manara dataset).

When we consider the Inverse Simpson Index, the results are exactly the same (the same 48 comparisons are significant), but when we consider the Shannon diversity index, we find a small difference. In this case the difference between strains resistant and susceptible to trimethoprim-sulfamethoxazole in the 2016 Yamaji dataset is not significant, but the same comparison in the Addams-Sapper dataset is.

### Regression analysis for the entire dataset

In the previous sections, we showed that diversity is typically lower for resistant strains than it is for susceptible strains. We noticed that while most comparisons were significantly different (resistant strains less diverse), this was not the case when resistance was very common. There are several reasons why we expected this to be the case, which we detailed in the introduction. Here we will use a linear model to determine whether there is a significant effect of how common resistance is on the diversity of resistant samples. For this analysis, we worked with the normalized Gini-Simpson Index values (see Methods).

We fitted a linear model to the normalized Gini-Simpson Index values, with the fraction of samples that are resistant (FracRes) and the Dataset as explanatory variables and found that indeed, diversity goes up with increasing fraction of resistant samples (FracRes), see figure 3.

**Figure 3.**
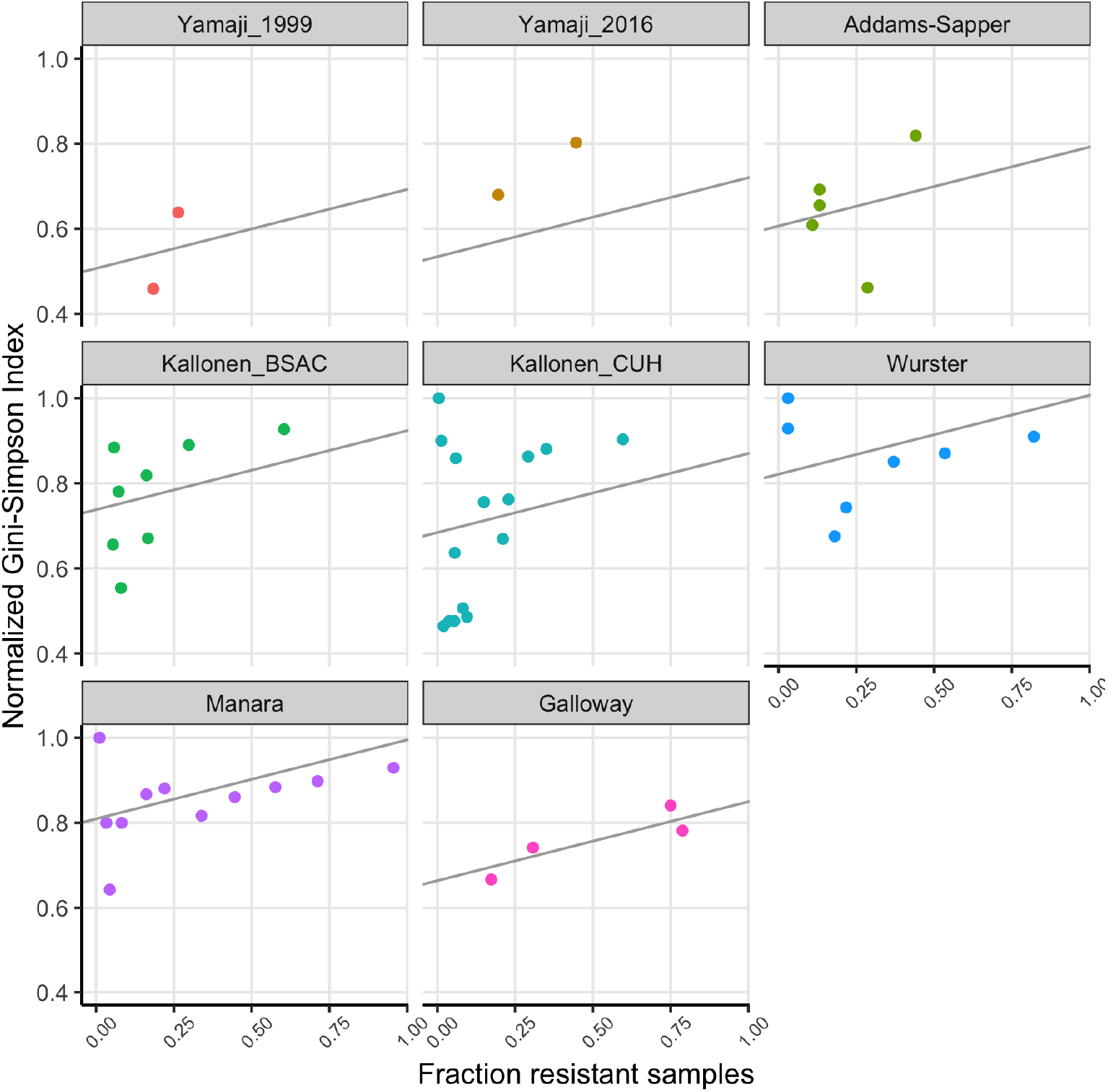
The fraction of samples that is resistant vs the normalized Gini-Simpson Index, with fitted linear model for each dataset.

## Discussion

In this study, we used existing clinical data on three different bacterial pathogens (*E. coli, S. aureus* and *E. faecium*) to determine whether drug resistant strains are less diverse than susceptible strains. Diversity here is considered at the between-host level and assessed by considering multi-locus sequence types. High diversity of resistant strains therefore means that different patients are infected with resistant strains of different multi-locus sequence types, whereas low diversity means that patients are infected with only a limited number of multi-locus sequence types represented among resistant strains.

We found that indeed, in all eight datasets, in most cases, resistant strains are less diverse than susceptible strains. All in all, in 51 of 59 comparisons, the resistant strains were significantly less diverse than susceptible strains (Figure 2). While this shows that resistant samples consist of fewer sequence types, the number of sequence types that carry resistance is larger than 1 in all cases. This shows that resistance has evolved or has been acquired (in the case of HGT) multiple times in every dataset and every specific antibiotic we consider.

Therefore, we can conclude that at the host population level, the evolution of antibiotic resistance typically occurs by multi-origin soft selective sweeps (Hermisson & Pennings, 2017; Messer & Petrov, 2013). This had previously been shown in a few other cases (Croucher et al., 2014; Nubel et al., 2008; Wilson et al., 2016), but we are not aware of another study that looked at multiple pathogens and multiple datasets.

Consistent with the theory of soft and hard selective sweeps (Hermisson & Pennings, 2017; Messer & Petrov, 2013), if resistance evolution or acquisition happens only rarely, we expect only few different sequence types to harbor drug resistance, which leads to low diversity of resistant strains. If resistance evolution or acquisition is common, for example, when horizontal gene transfer happens a lot, we expect many different sequence types to harbor drug resistance, leading to high diversity. In cases where HGT plays a role, we could also expect that when resistance becomes common, it could be transferred more easily to other strains within the same species, which will increase the sequence type diversity of resistant strains. This could be considered to be similar to when a partial selective sweep signature is lost over time due to recombination. There are thus multiple possible models that could lead to the observation of more diversity for more common resistances.

Consistent with these ideas, in the 8 comparisons where resistant samples were not significantly less diverse than susceptible strains, resistance was very common, with frequencies ranging from 44% to 96% of all samples being resistant. This result let us consider whether diversity of the resistant strains was generally affected by how common resistance is. We found that indeed, there was a significant effect of the fraction of samples that was resistant in a dataset for a given antibiotic and the diversity of these resistant strains (Figure 3).

The Yamaji datasets came from the same location (Northern California) but 17 years apart. In the first dataset (Yamaji 1999), trimethoprim-sulfamethoxazole-resistant and ampicillin-resistant strains are significantly less diverse than their susceptible counterparts. 17 years later (Yamaji 2016), diversity of the resistant strains has gone up in both cases. In addition, for ampicillin, the frequency of resistance has gone up (Figure 2A) and there is no longer a difference in diversity between resistant and susceptible strains. While this is a small dataset and the only case where we have longitudinal data, it suggests that over time, drug resistant strains can become both more common and more diverse.

Limitations of this study is that we had only eight datasets to work with. In some cases, the same genetic mechanism leads to resistance to multiple drugs (e.g., a plasmid with several resistance genes), but we did not take that into account in our analysis. This means that some of the 59 comparisons we tested should not be considered independent comparisons. However, because our results are quite similar between the different datasets and even different species of bacteria, we believe that our findings are quite general. Another statistical issue is the fact that some studies listed a small number of sequence types as “other” or “minor”. We removed these sequence types from our analysis. For most of the datasets, this is not an issue, or a very small issue (1 sample out of 53 for the Galloway dataset, 11 samples out of 184 for the Manara dataset), but for the Yamaji datasets, it affects a large number of samples (100 out of 224 (44%) in Yamaji 1999 and 85 out of 233 (36%) for Yamaji 2016). It is unfortunate that data is not always completely reported. Many other datasets we considered did not include patient-level sequence type data at all. We are hopeful that the recent push for data sharing will make this type of analysis more feasible in the future. We also expect that in the future, a similar study will be possible with genomic data instead of the limited sequence type data we used here.

With this study, we aim to bridge the world of population genetics, evolution and selective sweeps with the study of clinical bacterial samples and antibiotic resistance. A lot of effort goes into fighting antibiotic resistant infections. We believe that a better understanding of the evolution and spread of antibiotic resistant strains will ultimately contribute to finding better ways to reduce the number of antibiotic resistant infections.

## Acknowledgements

We would like to thank the SF BUILD writing retreat (August 15-18, 2022 Westerbeke Ranch Conference), we thank Dr Alison Feder and Dr Nandita Garud for discussions. AP and PSP were supported by NIH grant R01AI134195. LBR was supported by NIH MARC: T34-GM008574 and NIH SFSU/UCSF MS Bridges to the Doctorate: T32-GM142515. LBR and AP were supported by the Genentech Foundation. CC was supported by NIH MBRS-RISE: R25-GM059298.

